# Vasopressin 1a receptor antagonist disrupts male-male affiliative relationships formed by triadic cohabitation in large-billed crows

**DOI:** 10.1101/2024.02.01.578519

**Authors:** Akiko Seguchi, Ei-Ichi Izawa

**Affiliations:** Department of Psychology, Keio University, Tokyo 108-8345, Japan; Art Future Research Field, Tokyo University of the Arts, Tokyo 110-8714, Japan

**Author notes:** Correspondence: Akiko Seguchi Mita 2-15-45, Minato-ku, Tokyo 108-8345, Japan.

## Abstract

Same-sex affiliative relationships are common in humans and some social animals, forming one of the bases of group living. The neuropeptide vasopressin (VP) and its receptors mediate these relationships and behaviours in mammals and birds with gregarious and colonial social structures. In some species, affiliative relationships between dominant and subordinate individuals can be maintained while still retaining strict dominance hierarchies where three or more individuals interact. However, it is unclear whether triadic interaction promotes these relationships, and whether the VP system is also involved in such affiliations due to the lack of suitable animal models and experimental settings. This study addresses these questions with two experiments. In Experiment 1, two-week cohabitation among three male crows facilitated affiliative relationships in particular dyads within each triad. In Experiment 2, vasopressin 1a receptor (V1aR) antagonism disrupted affiliative behaviours and led to the resurgence of agonistic behaviours in affiliated males but not in unaffiliated ones by peripherally administering a V1aR antagonist. These findings suggest that the VP system might universally mediate same-sex affiliative relationships, despite differences in inherent aggression levels among individuals. The triadic cohabitation paradigm established here could advance our understanding of animal societies and be applied across various species, sexes, and social structures.

**Impact statement:** This study provides evidence that a triadic interactive environment facilitates the formation of affiliative relationship between specific dominant and subordinate males in large-billed crows, with crucial involvement of vasopressin 1a receptor in maintaining this affiliation.

## Introduction

Affiliative relationships, such as parent-offspring bonds (Clutton–Brock, 1991; Royle et al., 2012), male-female pair-bonds (Black, 1996; Kleiman, 1977; Bales et al., 2021), and same-sex platonic bonds (Brent et al., 2014), are beneficial for individuals living in groups as they increase the fitness of resource acquisition and, ultimately, reproductive success (Ziegler & Crockford, 2017). Data on the functions and neuroendocrine system of same-sex affiliative relationships, characterised by the exchange of non-sexual affiliative behaviours between same-sex individuals, including social grooming (allogrooming in mammals and allopreening in birds), huddling, spending time together, and sharing food, has predominantly have been obtained from studies on mammals and birds (Brent et al., 2014; Ziegler & Crockford, 2017; Beery, 2019). These relationships are not limited to one sex or kinship, as they have been observed in males and females, as well as in kin and non-kin, both in captivity and wild conditions (Ziegler & Crockford, 2017). Same-sex affiliative relationships are also frequent in humans, existing throughout most of our history and in many societies and cultures (Brent et al., 2014).

The neural mechanisms underlying affiliative relationships might have been co-opted from parent-offspring bonds into more flexible bond structures, such as pair bonds and same-sex affiliative relationships (Broad et al., 2006; Crespi, 2016; Numan & Young, 2016). The role of neuropeptides of the arginine vasopressin family (arginine vasotocin, AVT, in non-mammals) and oxytocin (OT) signals mediated V1aR has been implicated in same-sex affiliative relationships. This has been suggested by previous studies specifically in males and females of gregarious birds and females of voles. These studies primarily used paradigms that investigated “gregariousness” in finches as a preference for the larger of two groups, and partner preference in voles as a choice of familiar over unfamiliar. Goodson and Wang (2006) found that VT neurons in the medial bed nucleus of the stria terminalis were activated, in terms of immediate early gene Fos expression, in response to the exposure of same-sex conspecifics through the wire barrier in males and females of gregarious species, but were not in asocial ones. The authors also indicated that the activation of VT neurons is specifically induced by social stimuli with a positive valence to enhance affiliative behaviour (Goodson & Wang, 2006). Antisense-based knockdown of VT production in the medial bed nucleus of the stria terminalis of male zebra finches, a gregarious species, increased aggressive behaviour and reduced courtship in colony housing situations (Kelly & Goodson, 2013). V1aR antagonism in the LS of male zebra finches reduces flock size in housing conditions (Kelly et al., 2011). A previous study showed that in female meadow voles (*Microtus pennsylvanicus*), a season-specific social species, the co-administration of OT and a vasopressin 1a receptor (V1aR) antagonist in the LS inhibited female-female partner preference for huddling behaviour with cagemates over unfamiliar individuals (Anacker et al., 2016). Research into the physiological underpinnings of same-sex affiliation in voles is extensive and includes studies on neurotransmitters beyond VP/OT, such as the possible relevance of estradiol in meadow voles (Beery et al., 2008) and dopamine signalling in prairie voles (*M. ochrogaster*) with female-female partner preference (Lee & Beery, 2021; Lee et al., 2023).

However, among same-sex affiliative relationships, there is a type where individuals selectively affiliate with specific individuals despite the presence of multiple partners. Particularly, adult males can simultaneously hold a position within strict dominance relationships and form affiliative relationships between specific dominant and subordinate individuals. Such relationships, including coalitions and alliances, have been observed in various animal species (including humans, *Homo sapiens* (Macfarlan et al., 2014; von Rueden et al., 2008; von Rueden & Jaeggi, 2016; Patton, 2005); chimpanzees, *Pan troglodytes* (Watts, 2002; Muller & Mitani, 2005; Gilby et al., 2013); baboons, *Papio* (Bercovitch, 1988; Noë, 1994); macaques, *Macaca radiata* (Silk, 1999); *M. fuscata* (Kutsukake & Hasegawa, 2005); cetaceans bottlenose dolphins, *Tursiops truncates* (Connor et al., 1992a; Connor et al., 1992b; Connor et al., 1999); spotted dolphins, *Stenella frontalis* (Green et al., 2015); sperm whales, *Physeter macrocephalus* (Kobayashi et al., 2020); carnivores like lions, *Panthera leo* (Bygott et al., 1979); cheetahs, *Acinonyx jubatus* (Caro, 1993; Caro, 1994); birds, corvids (Fraser & Bugnyar, 2012). Most of these studies indicated that male-male affiliative relationships were formed in social interactions involving multiple participants, typically two individuals vs. one individual, in both the wild and captivity. Theoretical research exploring the social factors of coalition formation predicted that social interactions among three individuals differing in “power” and “activity” should be the minimal group size to produce coalition/alliance (Caplow, 1956; Mesterton-Gibbons et al., 2011; Koykka & Wild, 2017). Several previous studies with human participants have examined the combinations of ranks that facilitate the formation of coalitions in game situations (Vinacke & Arkoff, 1957; Kelley & Arrowood, 1960). If replicated in experimental research on social animals, the formation of affiliative relationships between males within these three-individual interactions can facilitate studies on the physiological basis of such relationships.

Crows and related species (i.e., *Corvus* spp) are an ideal model to investigate the ecological factors and physiological basis of male-male affiliative relationships. Non-breeding crows form groups during foraging, roosting, and socialising, with group sizes ranging from a few to several hundred or even thousands of birds and with the composition of group members changing during the day and across contexts (Boucherie et al., 2019; Izawa, 2011; Loretto et al., 2017; Uhl et al., 2019). Despite the high degree of fission-fusion dynamics, temporarily stable sub-groups can be observed. For example, certain birds use the same food source or roost repeatedly over consecutive weeks or years (Braun & Bugnyar, 2012). Additionally, individuals form multiple highly differentiated affiliative and agonistic relationships with others that comprise various qualities (Fraser & Bugnyar, 2010a). Males form and maintain stable dyadic dominance relationships and, in a group setting, a strict linear hierarchy, whereas those of females are less robust (Fraser & Bugnyar, 2010a; Izawa & Watanabe, 2008; Nishizawa et al., 2011; Ode et al., 2015; Takeda et al., 2022).

In both captive and wild populations, affiliative relationships are formed between unpaired adults and unmatured juveniles by exchanging affiliative behaviour such as allopreening, food giving, and conflict aiding (Bugnyar, 2012; de Kort et al., 2006; Emery et al., 2007a; Scheid et al., 2008; von Bayern et al., 2007; Picard et al., 2020). Male-male and male-female dyads tend to form stronger and more stable affiliative relationships than female-female dyads (Fraser & Bugnyar, 2010a; Miyazawa et al., 2020). The formation of affiliative relationships between individuals increases the dominance rank and likelihood of success in competing for food in captive groups of rooks (*Corvus frugilegus*) (Emery et al., 2007b) and wild common ravens (*C. corax*) (Braun & Bugnyar, 2012; Braun et al., 2012). These findings suggest an advantage of the affiliative relationship in group lives. Ravens and rooks reconcile with and console valuable partners (Fraser & Bugnyar, 2010b; Fraser & Bugnyar, 2011; Seed et al., 2007), remember former group mates and their relationships with them over years (Boeckle & Bugnyar, 2012), understand third-party relations (Massen et al., 2014a), and prevent others from forming too strong alliances (Massen et al., 2014b). Moreover, ravens cooperate in the wild; for example, they support each other in conflicts (Fraser & Bugnyar, 2012), hunt cooperatively (Hendricks & Schlang, 1998) and cooperatively chase away larger predators or dominant male-female pairs from prey items (Heinrich & Marzluff, 1991; Marzluff et al., 1996). Like other *Corvus* species, non-breeders of large-billed crows (*C. macrorhynchos*) form fission–fusion societies, with flexible changes in the social structure depending on the time of day, seasons, and their age (Izawa, 2011; Islam et al., 2010; Kuroda, 1990). In captive non-breeder groups, males strictly form a linear dominance hierarchy (Izawa & Watanabe, 2008; Ode et al., 2015) and allopreening often occurs between males (Miyazawa et al., 2020). This allopreening is primarily initiated by dominants but also occurs from subordinates (Miyazawa et al., 2020).

A triadic interactive scenario is an ecological factor that facilitates the formation of affiliative relationships between dominant and subordinate males. Previous studies have suggested the involvement of V1aR in same-sex affiliative relationships; therefore, in this study, we hypothesised that cohabitation of three male non-breeder large-billed crows could facilitate the formation of affiliative relationships between specific dominant and subordinate individuals and that V1aR is involved in the maintenance of these relationships. To this end, we conducted two experiments to test our hypotheses.

In Experiment 1, we introduced three non-breeder males of captive large-billed crows, which had formed dominance relationships but not affiliative relationships prior to this study, to a triadic cohabitation in an aviary for two weeks. We examined the formation of an affiliative relationship in one of the three possible dyads by comparing the occurrence of reciprocal allopreening before and after cohabitation. As we predicted, the two-week triadic cohabitation facilitated the formation of an affiliative relationship exclusively between two males in the triad. The two-week triadic cohabitation paradigm developed in this study enabled us to investigate the physiological mechanisms of male-male affiliative relationships in a laboratory setting.

In Experiment 2, we investigated the effects of peripheral administration of a V1aR antagonist on the social behaviours of males with affiliative relationships formed via triadic cohabitation. Our prediction was that blocking V1aR would reduce allopreening and increase aggressive and submissive behaviours among dominant and subordinate males with affiliative relationships but not among those without such relationships. We discovered that administering a V1aR antagonist disrupted the affiliative relationships between males, leading to more aggressive and submissive behaviours from dominants and subordinates. However, it had no effect on the social behaviour of males without affiliative relationships. In conclusion, our study suggests that living in a group of three promotes the formation of close relationships between males and that V1aR is involved in maintaining these relationships by promoting allopreening and inhibiting aggressive and submissive behaviours based on dominance.

## Results

### Experiment 1: The triadic cohabitation facilitated the formation of an affiliative relationship between two particular males

To determine whether triadic cohabitation among males promotes the formation of affiliative relationships between specific dyads, we introduced eight groups of three male crows, each already having established dominance relationships but no affiliative relationships, into an aviary for two weeks (Figure 1). We compared social behaviour between males before and after cohabitation to evaluate the formation of affiliative relationships characterised by reciprocal allopreening exclusively between two of three males in each triad.

**Figure 1.**
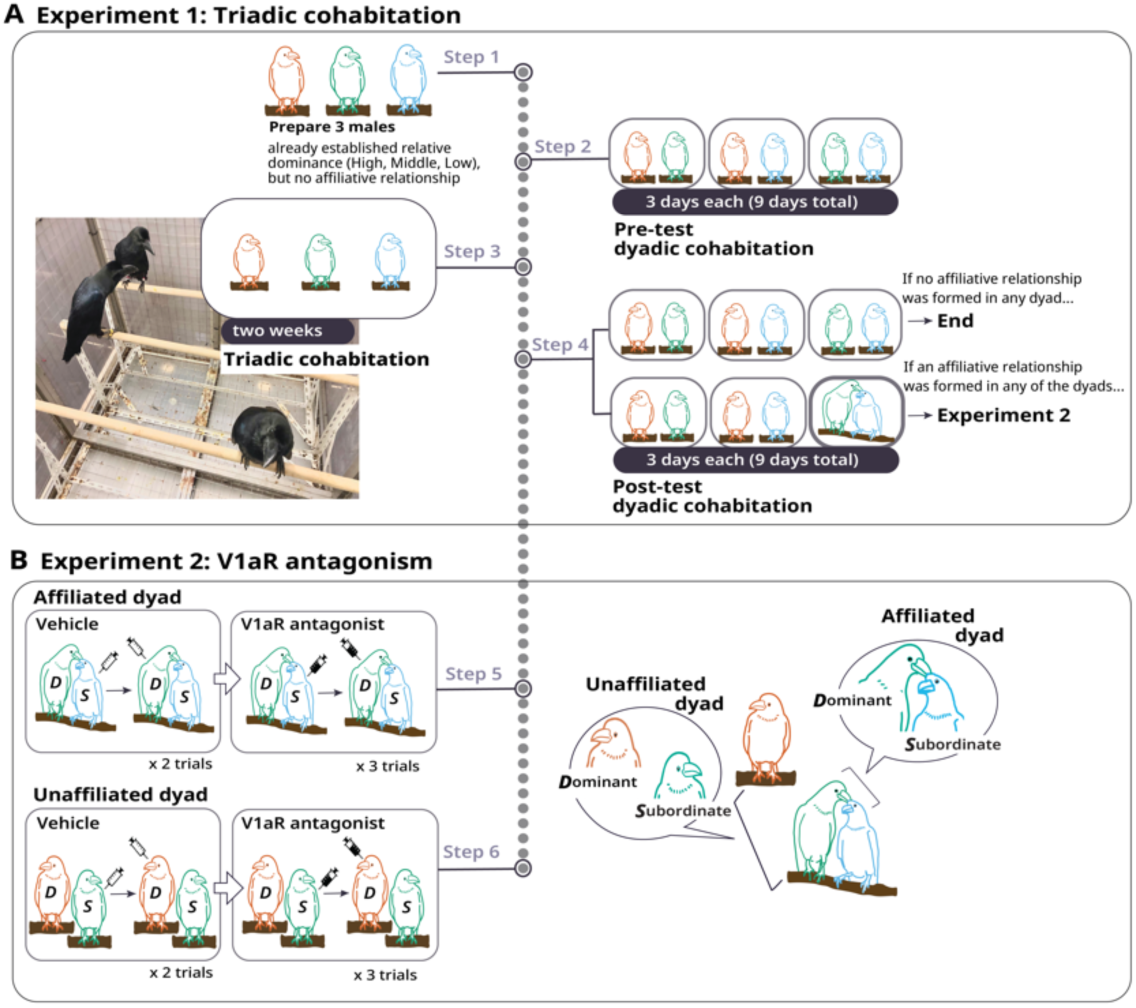
Experimental design. (A) Experiment 1: Triadic cohabitation and formation of affiliative relationships. Step 1: three male crows with already established relative dominance relationships through each dyadic encounter but no affiliative relationships were introduced into an aviary for two weeks. These three crows are referred to as high (orange), middle (green), and low (blue) based on their relative dominance. Step 2: pre-test dyadic cohabitation for 3 days to ensure any affiliative relationships noted were not due to prior dyadic interactions. Step 3: triadic cohabitation for two weeks to observe the formation of affiliative relationships, characterised by reciprocal allopreening between two specific crows in each triad. Step 4: post-test dyadic cohabitation for 3 days to confirm the persistence of these relationships. If no affiliative relationship was formed in any dyad, that triad would not proceed to Experiment 2, the behavioural pharmacological experiment. (B) Experiment 2: Effects of V1a receptor antagonism on affiliative relationships between dominant and subordinate individuals. Steps 5 and 6: V1aR antagonists and control vehicle solutions were administered peripherally to one of the two individuals with established affiliative or unaffiliative relationships from Experiment 1, prior to their dyadic interaction. In each trial, the drug to either the dominant or subordinate individual, and free interactions between the two were compared pre- and post-administration. After completing all vehicle trials, the antagonist was administered following the same protocol. Allopreening, aggressive behaviours and submissive vocalisations, were coded for analysis to determine the influence of the V1aR antagonist on the maintenance of affiliative behaviours and the emergence of aggression or submission.

Before this experiment, dominance relationships of all dyads used in this study were assessed using a social dyadic encounter paradigm from previous studies (Nishizawa et al., 2011; Takeda et al., 2022). Briefly, two birds were introduced into an aviary to freely interact for 5 min in three trials within a 2-day inter-trial interval. In each trial, the individual exhibiting submissive behaviours in response to aggressive behaviours from the opponent was identified as the loser, while the other as the winner. Within the dyads, the individual who won all three trials was defined as dominant, while the others as subordinates. All 26 dyads comprising the eight triads tested in this study showed a clear asymmetry in win/loss numbers, indicating established dominance relationships in all the dyads (Table 1). In these dyadic encounter trials, we also measured the number of allopreening and confirmed none occurred (Figure 2, Figure 2—source data 1). These results indicated that dominance, but no affiliative, relationships were established in all the dyads before Experiment 1.

**Figure 2.**
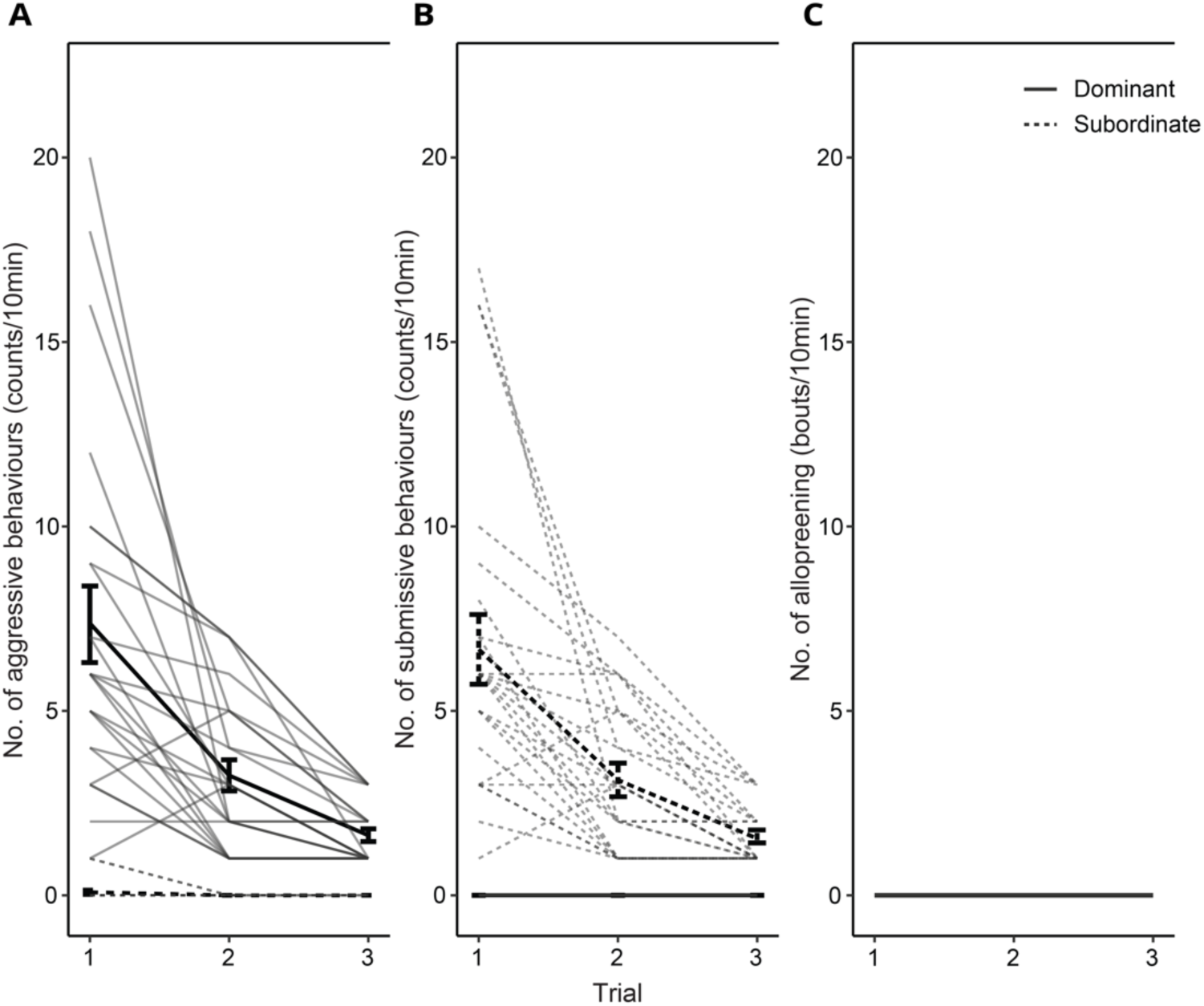
Determination of relative dominance in dyads. The numbers of aggressive (A), submissive (B) and allopreening behaviours (C) across the three trials of dyadic social encounters. The numbers of aggressive behaviours (jab, peck, aggressive vocalisation and displacing approach) by the dominant (thick solid black line) and submissive behaviours (submissive begging vocalisation and avoidance) by the subordinates (thick dotted black line) decreased through the three encounters, suggesting the stable formation of dyadic dominance relationships. Allopreening behaviours were not observed in any dyads. Grey-coloured solid and dotted lines indicate dominant and subordinate individual data, respectively. Data are represented with mean ± s.e.

**Table 1.**
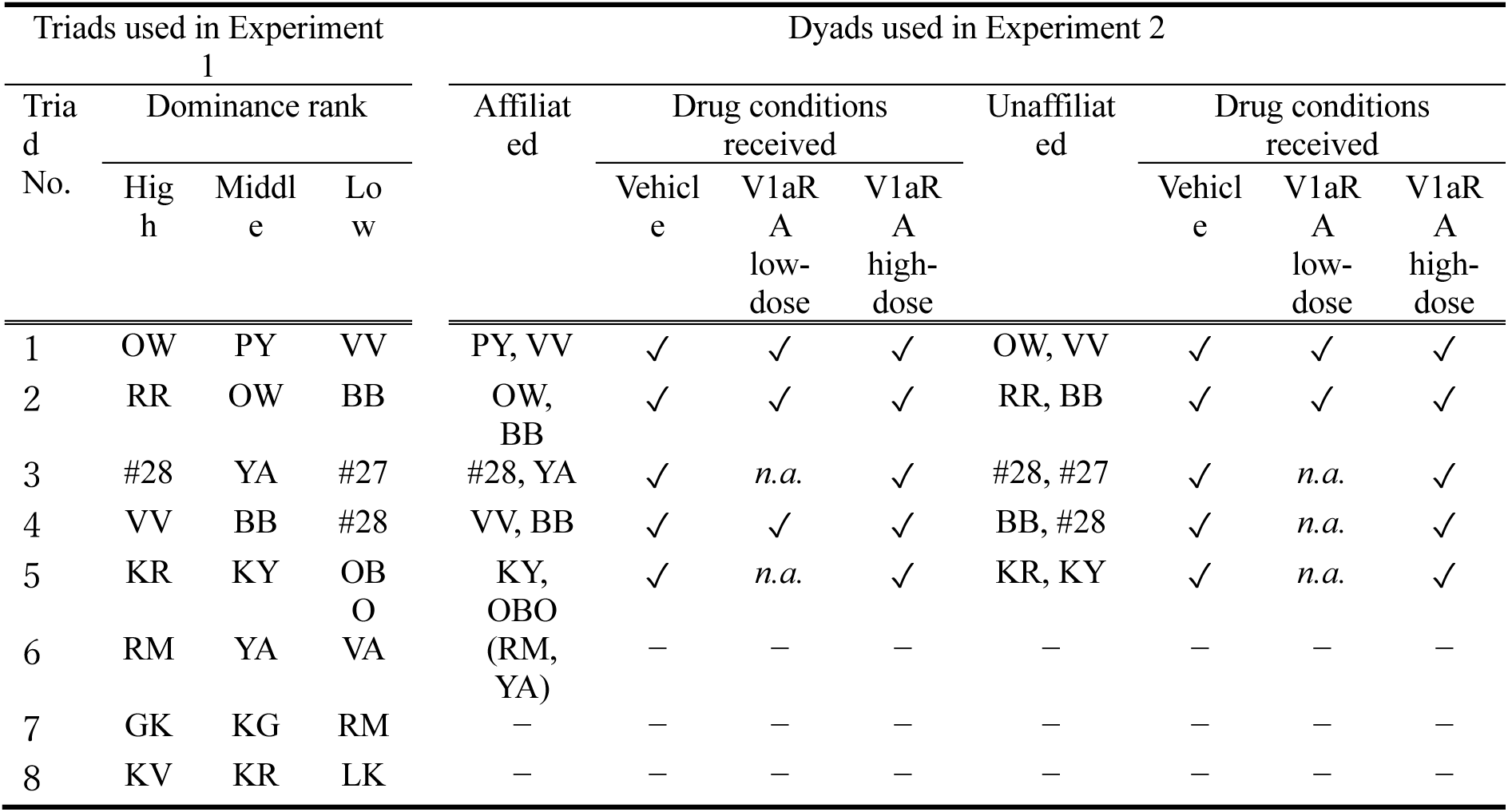
Individuals used in Experiments 1 and 2. Birds OW, YA, VV, BB, #28, and KR are included in two different triads. Triad #6 (RM, YA, VA) did not receive Experiment 2 (see text for the details). *n.a.* indicates data is not available.

Prior to triadic cohabitation (the pre-test), we housed all potential dyads together in the experimental aviary for three days to ensure that any affiliative relationships observed were not due to prior dyadic interactions. This was followed by a 2-week triadic cohabitation to determine whether such interactive situations would encourage affiliative relationships, evidenced by increased reciprocal allopreening between two specific birds in each triad. To confirm the persistence of these relationships, a follow-up dyadic cohabitation (the post-test) was conducted after the initial 2-week period.

Affiliative relationships were defined by the exchange of reciprocal allopreening between two birds, with each individual initiating allopreening at least 10 bouts per 30 min in any of the three 3-day blocks (early, middle and last) during the triadic cohabitation. Dyads not reaching this threshold were considered unaffiliated. Discriminant analysis, with leave-one-out cross-validation, was employed to differentiate the numbers of dominant-initiated and subordinate-initiated allopreening between the pre-test and post-test and to validate the exclusivity of the affiliative relationships formed during triadic cohabitation.

In six of the eight triads, the 2-week cohabitation resulted in the formation of an affiliative relationship exclusively in one dyad but not in the other dyads (Figure 3A, Figure 3—source data 1, Video 1). A specific dyad in each triad (#1–#6), which showed no allopreening in the pre-test, exhibited an increase in allopreening through the 2-week triadic cohabitation. In the post-test of these six dyads, allopreening was initiated by both dominants and subordinates, whereas no reciprocal allopreening occurred in the pre-test, and no such allopreening emerged in any other dyads in either test (Figure 3A, Figure 3—source data 1). Discriminant analysis based on the number of allopreening initiated by dominants and subordinates within the dyads yielded two discriminant functions. Using the leave-one-out method, the two discriminant functions distinguished the affiliated dyads in the post-test from themselves in the pre-test and from the other unaffiliated dyads with an 83.3% correct rate (Figure 3B, Figure 3—source data 2, i.e., five of six dyads were classified as affiliated dyads; see the outputs for Table 2). These results indicate that the 2-week triadic cohabitation facilitates the formation of affiliative relationships exclusively between two of the three birds, characterised by reciprocal allopreening.

**Figure 3.**
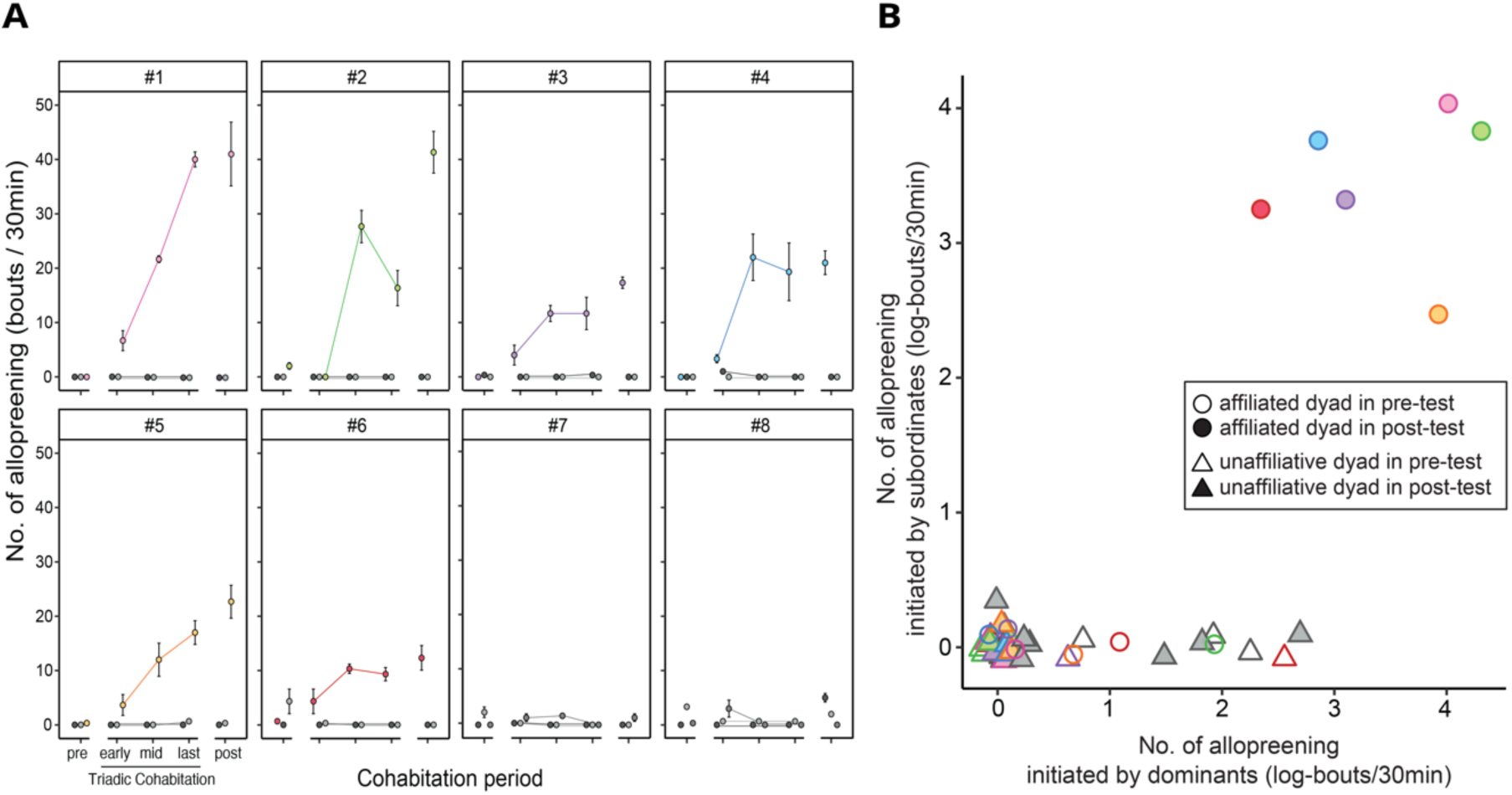
Triadic cohabitation facilitated the formation of an affiliative relationship between two particular males. (A) Total number of allopreening behaviours in each dyad across the experimental periods: pre-tests, 2-week triadic cohabitation (early, middle, and last), and post-tests. Data plots are presented as mean ± 1 s.e. Panels indicate triads (#1−#8). Bright-coloured plots (pink, green, purple, blue, orange and red) in the #1−#6 triads represent dyads that showed the formation of affiliated relationships. Darker plots denote the dyads that did not form affiliative relationships. (B) Scatter plots of allopreening behaviours initiated by dominants (x-axis) versus subordinates (y-axis) of dyads in the pre-tests (opened) and the post-tests (shaded). Coloured plots represent dyads wherein affiliative relationships were formed. Allopreening occurred exclusively in the coloured dyads after the cohabitation (i.e., post-test). The colours correspond to the dyad in Figure 3A. Data are presented as the mean with natural log-transformed values.

**Table 2.**
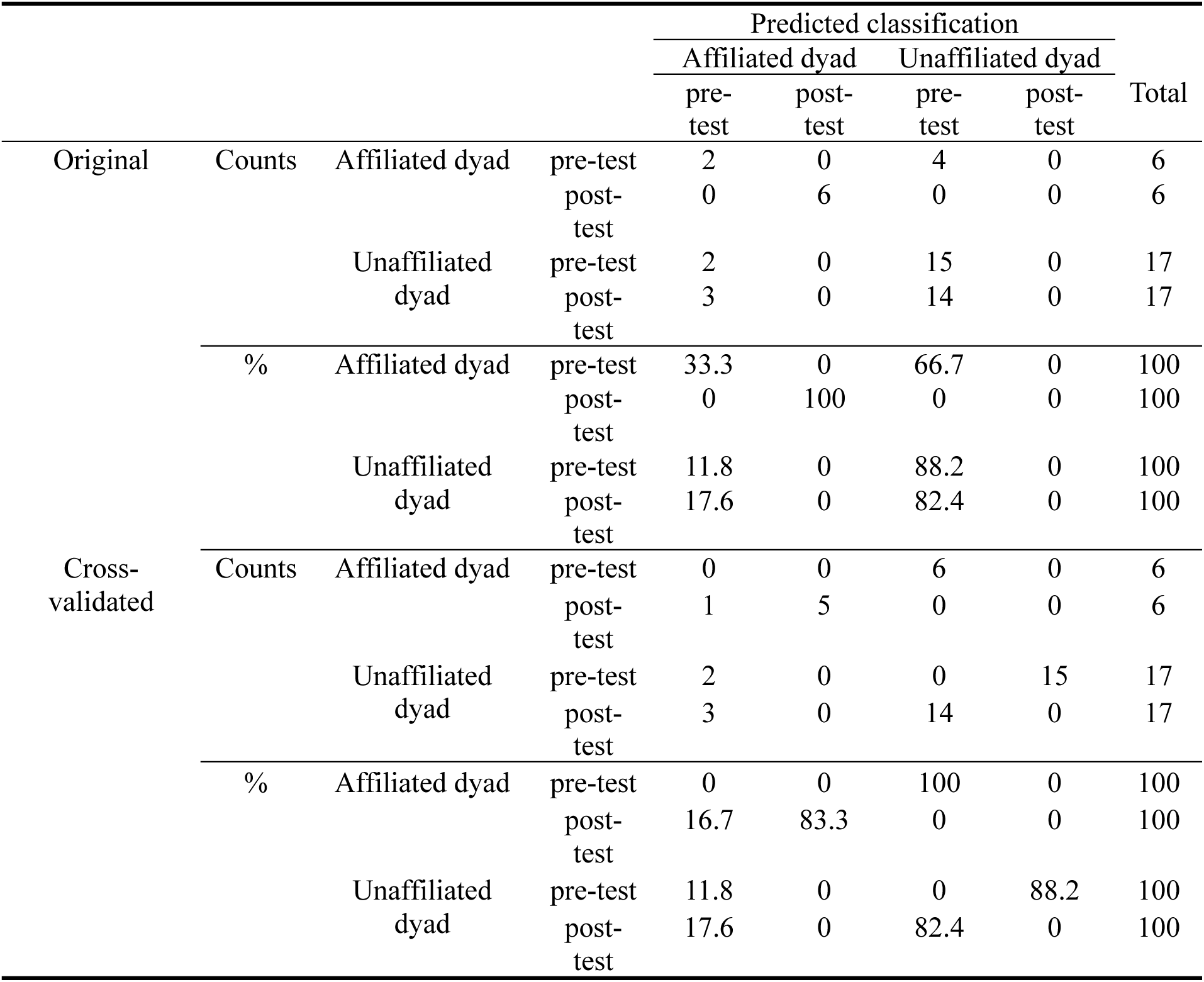
Outputs of discriminant analysis.

Due to a physical injury in one bird, five of the six dyads forming an affiliative relationship were used for the test in Experiment 2 (Table 1).

### Experiment 2: V1a receptor antagonism disrupted dominant-subordinate affiliative relationships

To elucidate the role of V1a-like receptors in the maintenance of male-male affiliative relationships, we systematically administered a V1aR antagonist to crows with pre-established affiliative or unaffiliative relationships from Experiment 1, aiming to assess its effects on their social interactions with both affiliative and unaffiliative counterparts (see Materials and Methods, Exp2). The antagonist was administered intramuscularly at both high and low doses, alongside a control vehicle solution. In each trial, the two individuals were first recorded interacting in the aviary for 30 minutes in the morning (pre-administration phase). Following this, either the dominant or subordinate crow received the antagonist or vehicle. One-hour post-administration, the crows were again observed for 30 minutes (post-administration phase). Social behaviours, including allopreening, aggressive behaviours, and submissive vocalizations were recorded via video and analysed subsequently. By comparing the frequency of these behaviours between the pre- and post-administration phases, this experiment aimed to clarify the impact of V1aR antagonism on the maintenance of established affiliative behaviours, as well as the potential induction of aggression and submission.

The administration of the V1aR antagonist significantly reduced allopreening in affiliated dyads, including both dominant and subordinate individuals (Figure 4, Figure 4-6—source data 1, Video 2). The Generalized Linear Mixed Model (GLMM) analysis revealed a significant interaction between drug condition and phase in both the low-dose and high-dose conditions (*V1aRA low-dose*, estimate ± s.e. = -2.231 ± 0.278, *p* < 0.001; *V1aRA high-dose*, estimate ± s.e. = -4.111 ± 0.592, *p* < 0.001; Table 3). A three-way interaction between dominance status, drug condition, and phase was significant only in the high-dose condition (estimate ± s.e. = 1.580 ± 0.638, *p* < 0.05; Table 3). Post-hoc pairwise comparisons further clarified that allopreening in affiliated dyads significantly decreased in the post-administration phase under both low-dose and high-dose conditions of the V1aR antagonist (*Pre vs. Post for V1aRA low-dose in dominants*, estimate ± s.e. = 1.798 ± 0.208, *p* < 0.001; *Pre vs. Post for V1aRA low-dose in subordinates*, estimate ± s.e. = 2.354 ± 0.254, *p* < 0.001; *Pre vs. Post for V1aRA high-dose in dominants*, estimate ± s.e. = 2.453 ± 0.213, *p* < 0.001; *Pre vs. Post for V1aRA high-dose in subordinates*, estimate ± s.e. = 4.234 ± 0.581, *p* < 0.001; Table 4). Additionally, post-hoc pairwise comparisons of drug conditions on allopreening in affiliated dyads during the post-administration phase showed that both low-dose and high-dose conditions significantly reduced allopreening compared to the vehicle condition in dominants (*Vehicle vs. V1aRA low-dose*, estimate ± s.e. = 2.047 ± 0.208, *p* < 0.001; *Vehicle vs. V1aRA high-dose*, estimate ± s.e. = 2.327 ± 0.217, *p* < 0.001; Table 5). No significant difference was observed between the low-dose and high-dose conditions in dominants (estimate ± s.e. = 0.280 ± 0.281, *p* = 0.579; Table 5). In subordinates, both low-dose and high-dose conditions also significantly reduced allopreening compared to the vehicle condition (*Vehicle vs. V1aRA low-dose*, estimate ± s.e. = 2.255 ± 0.258, *p* < 0.001; *Vehicle vs. V1aRA high-dose*, estimate ± s.e. = 4.152 ± 0.583, *p* < 0.001; Table 5). Notably, a significant difference was found between the low-dose and high-dose conditions in subordinates, with the high-dose condition showing a more pronounced reduction in allopreening (*V1aRA low-dose vs. V1aRA high-dose*, estimate ± s.e. = 1.897 ± 0.626, *p* < 0.01; Table 5). These findings indicate that V1aR antagonism reduces allopreening behaviour in both dominants and subordinates, with the effect being more pronounced in subordinates under the high-dose condition. No significant effects of V1aR antagonist administration on allopreening were observed in unaffiliated dyads (Figure 4). GLMM analysis was not conducted on unaffiliated dyads, as no allopreening was recorded under any condition or with any bird.

**Figure 4.**
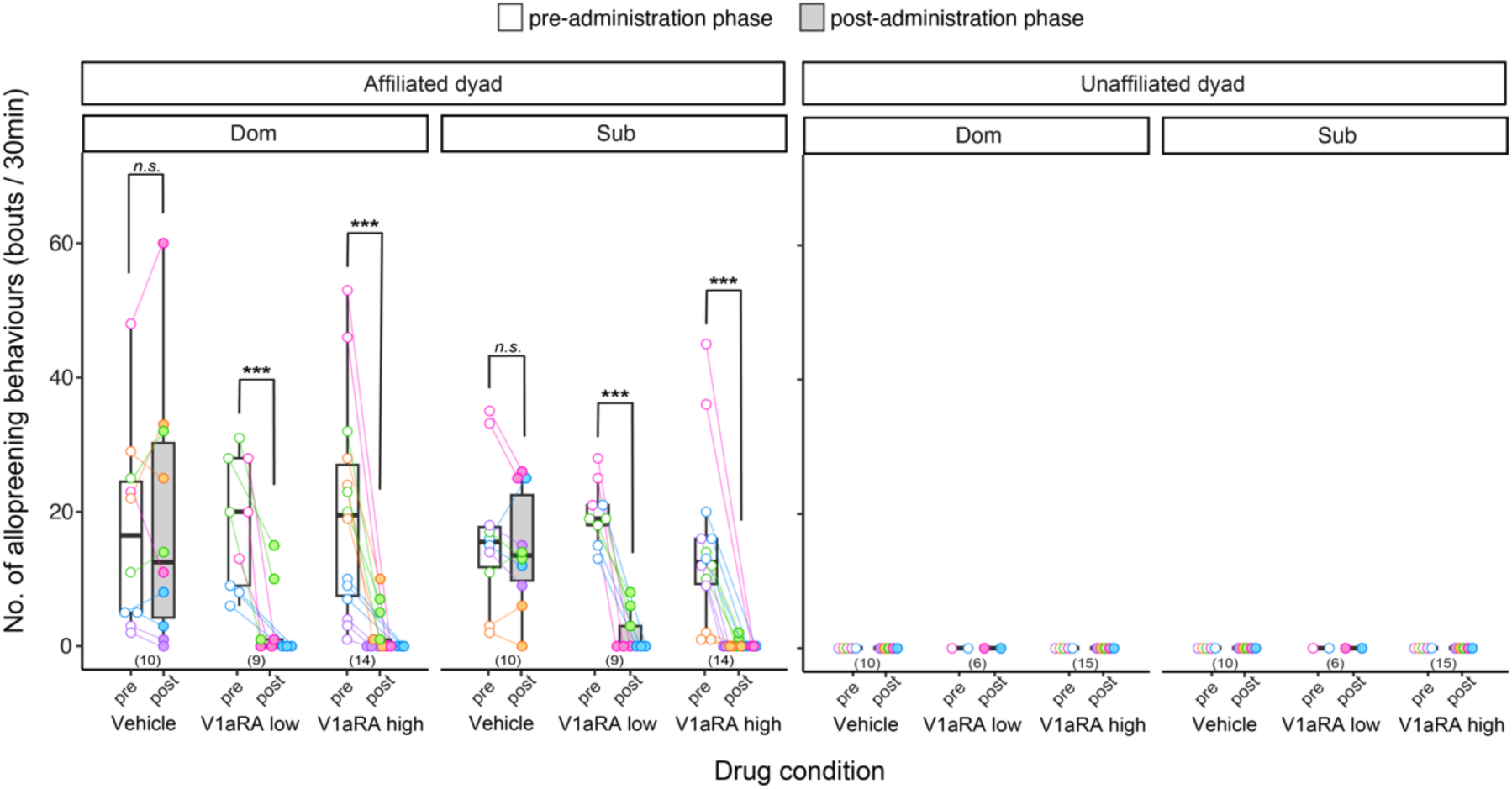
Effects of V1a receptor antagonist (V1aRA) on the number of allopreening behaviours in affiliated and unaffiliated dyads. In affiliated dyads, allopreening behaviours initiated by both dominant (DOM, left panel) and subordinate (SUB, right panel) crows exhibited a significant decrease in both the *high-dose* and *low-dose* V1aRA conditions post-administration, compared to pre-administration. Conversely, in the vehicle condition, no significant difference was observed in allopreening frequency between pre- and post-administration phases. In contrast, within unaffiliated dyads, V1aRA administration exerted no effect on allopreening for either the dominant (DOM, left panel) or subordinate (SUB, right panel) individuals. The plots illustrate individual data for each trial. The upper, middle, and lower bounds of the boxes represent the 75^th^, 50^th^, and 25^th^ percentiles, respectively, with whiskers extending to the maximum and minimum values. Numbers in the brackets below the boxes denote the number of samples (no. of birds × no. of trials). Asterisks denote statistically significant differences (Wald *χ^2^*statistics, *** *p* < 0.001). The colours correspond to the affiliated dyads shown in Figure 3.

**Table 3.** Outputs of GLMMs for allopreening behaviour in affiliated dyads.

| variables | estimates | s.e. | *z* -value | *p* |
| --- | --- | --- | --- | --- |
| intercept | 2.672 | 0.238 | 11.225 | < 0.001 |
| Subject dominance status<sup>a</sup> | 0.053 | 0.109 | 0.491 | 0.624 |
| Drug condition<sup>b</sup> |  |  |  |  |
| V1aRA low-dose | -0.024 | 0.112 | -0.211 | 0.833 |
| V1aRA high-dose | -0.041 | 0.106 | -0.384 | 0.701 |
| Phase<sup>c</sup> | -0.123 | 0.114 | -1.081 | 0.280 |
| Subject dominance status<sup>a</sup>×Drug condition<sup>b</sup> |  |  |  |  |
| ×V1aRA low-dose | -0.147 | 0.154 | -0.958 | 0.338 |
| ×V1aRA high-dose | 0.245 | 0.142 | 1.722 | 0.085 |
| Subject dominance status<sup>a</sup>×Phase<sup>c</sup> | 0.201 | 0.155 | 1.295 | 0.195 |
| Drug condition<sup>b</sup>×Phase<sup>c</sup> |  |  |  |  |
| ×V1aRA low-dose | -2.231 | 0.278 | -8.024 | < 0.001 |
| ×V1aRA high-dose | -4.111 | 0.592 | -6.944 | < 0.001 |
| Subject dominance status<sup>a</sup>×Drug condition<sup>b</sup>×Phase<sup>c</sup> |  |  |  |  |
| ×V1aRA low-dose | 0.355 | 0.363 | 0.980 | 0.327 |
| ×V1aRA high-dose | 1.580 | 0.638 | 2.477 | < 0.05 |

Italicized indicates statistically significant results.
<sup>a</sup> Subordinate was set to 0.
<sup>b</sup> Vehicle was set to 0.
<sup>c</sup> Pre was set to 0.

**Table 4.** Post-hoc pairwise comparisons of pre- and post-administration on allopreening behaviour by drug conditions and subject dominance status.

| Drug condition | Subject dominance status | Contrast | estimate | s.e. | z-ratio | *p* |
| --- | --- | --- | --- | --- | --- | --- |
| Vehicle | Dominant | Pre vs. Post | -0.078 | 0.105 | -0.738 | 0.460 |
|  | Subordinate | Pre vs. Post | 0.123 | 0.114 | 1.081 | 0.280 |
| V1aRA low-dose | Dominant | Pre vs. Post | 1.798 | 0.208 | 8.658 | < 0.001 |
|  | Subordinate | Pre vs. Post | 2.354 | 0.254 | 9.282 | < 0.001 |
| V1aRA high-dose | Dominant | Pre vs. Post | 2.453 | 0.213 | 11.543 | < 0.001 |
|  | Subordinate | Pre vs. Post | 4.234 | 0.581 | 7.288 | < 0.001 |

Italicized indicates statistically significant results.

**Table 5.** Post-hoc pairwise comparisons of drug conditions on allopreening behaviour by subject dominance status in the post-administration phase.

| Subject dominance status | Contrast | estimate | s.e. | z-ratio | *p* |
| --- | --- | --- | --- | --- | --- |
| Dominant | Vehicle vs. V1aRA low-dose | 2.047 | 0.208 | 9.840 | < 0.001 |
|  | Vehicle vs. V1aRA high-dose | 2.327 | 0.217 | 10.704 | < 0.001 |
|  | V1aRA low-dose vs. V1aRA high-dose | 0.280 | 0.281 | 0.996 | 0.579 |
| Subordinate | Vehicle vs. V1aRA low-dose | 2.255 | 0.258 | 8.739 | < 0.001 |
|  | Vehicle vs. V1aRA high-dose | 4.152 | 0.583 | 7.121 | < 0.001 |
|  | V1aRA low-dose vs. V1aRA high-dose | 1.897 | 0.626 | 3.030 | < 0.01 |

Italicized indicates statistically significant results.

The administration of a high dose of the V1aR antagonist increased aggression from dominant individuals toward subordinates but did not enhance aggression from subordinates toward dominants in affiliated dyads (Figure 5, Figure 4-6—source data 1, Video 2). GLMM analysis revealed a significant interaction between drug condition and phase under the high-dose V1aR antagonist condition (estimate ± s.e. = 3.114 ± 1.065, *p* < 0.01; Table 6). Post-hoc pairwise comparisons further clarified that aggressive behaviours in affiliated dyads significantly increased during the post-administration phase under the high-dose condition of the V1aR antagonist (*Pre vs. Post*, estimate ± s.e. = -3.110 ± 0.705, *p* < 0.001; Table 7). Moreover, post-hoc comparisons of drug conditions on aggressive behaviours in affiliated dyads during the post-administration phase revealed that the high-dose condition significantly increased aggression compared to both the vehicle and low-dose conditions (*Vehicle vs. V1aRA high-dose*, estimate ± s.e. = -2.643 ± 0.584, *p* < 0.001; *V1aRA low-dose vs. V1aRA high-dose*, estimate ± s.e. = -2.281 ± 0.590, *p* < 0.001; Table 8). These findings indicate a substantial increase in aggression from dominant individuals following the administration of a high dose of the V1aR antagonist. No similar increase in aggression was observed in subordinate individuals (see Figure 5; GLMM analysis was not conducted due to the absence of aggression under any condition or in any bird). Furthermore, the V1aR antagonist did not enhance aggression in unaffiliated dyads (Figure 5 and Table 9).

**Figure 5.**
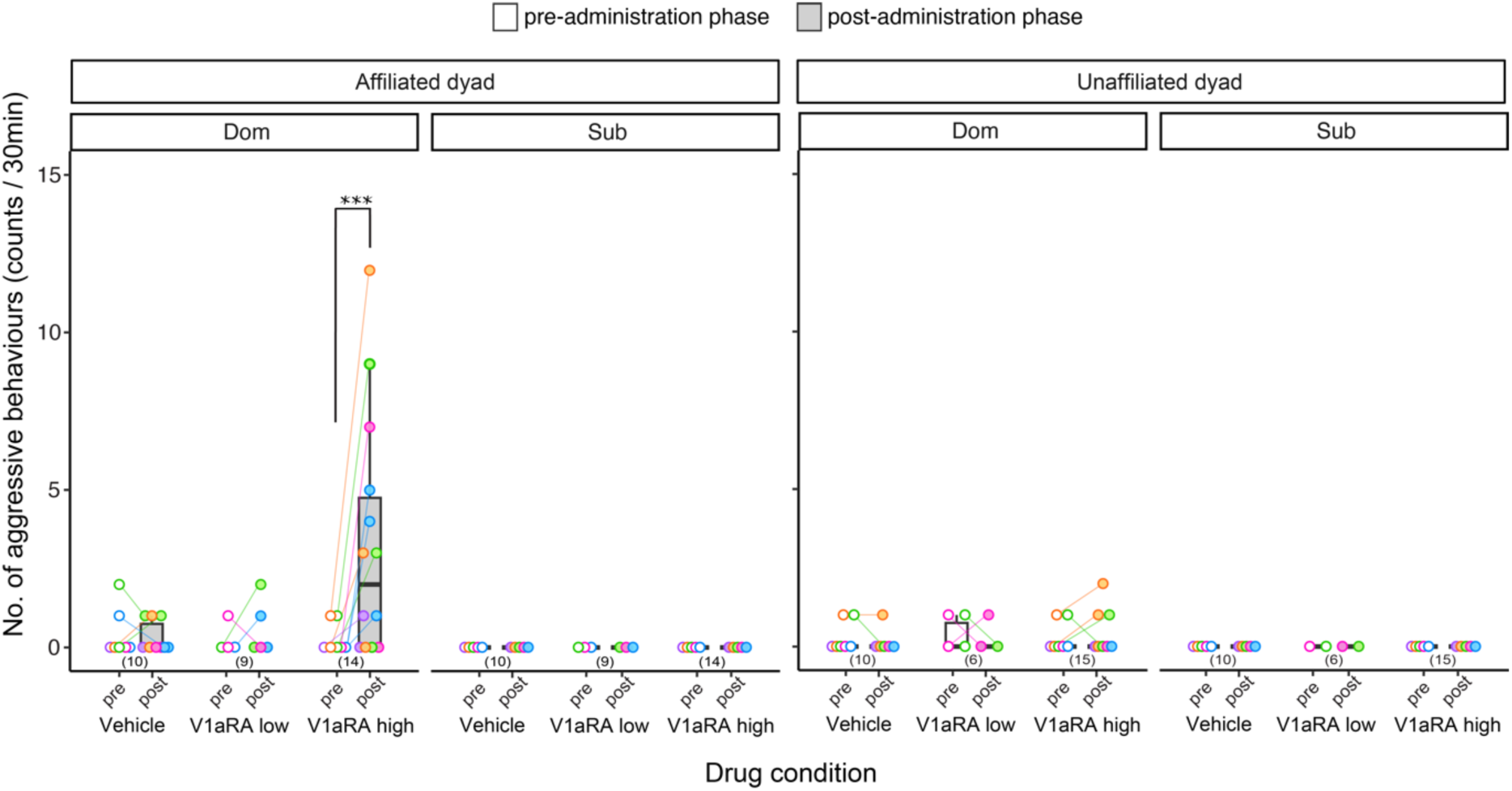
Effects of V1aRA on the number of aggressive behaviours in affiliated and unaffiliated dyads. In affiliated dyads, aggressive behaviour initiated by the dominant individual (DOM, left panel) exhibited a significant increase following administration of the *high-dose* V1aRA, as compared to pre-administration phase. In contrast, no such increase was observed in the low-dose or vehicle conditions. Additionally, this elevation in aggression was not observed in any condition for the subordinate individual (SUB, right panel). Furthermore, in unaffiliated dyads, no significant effects were detected in either the dominant or subordinate individuals under any of the administered conditions. The plots illustrate individual data for each trial. The upper, middle, and lower bounds of the boxes represent the 75^th^, 50^th^, and 25^th^ percentiles, respectively, with whiskers extending to the maximum and minimum values. Numbers in the brackets below the boxes denote the number of samples (no. of birds × no. of trials). Asterisks denote statistically significant differences (Wald *χ^2^*statistics, *** *p* < 0.001). The colours correspond to the affiliated dyads shown in Figure 3.

**Table 6.**
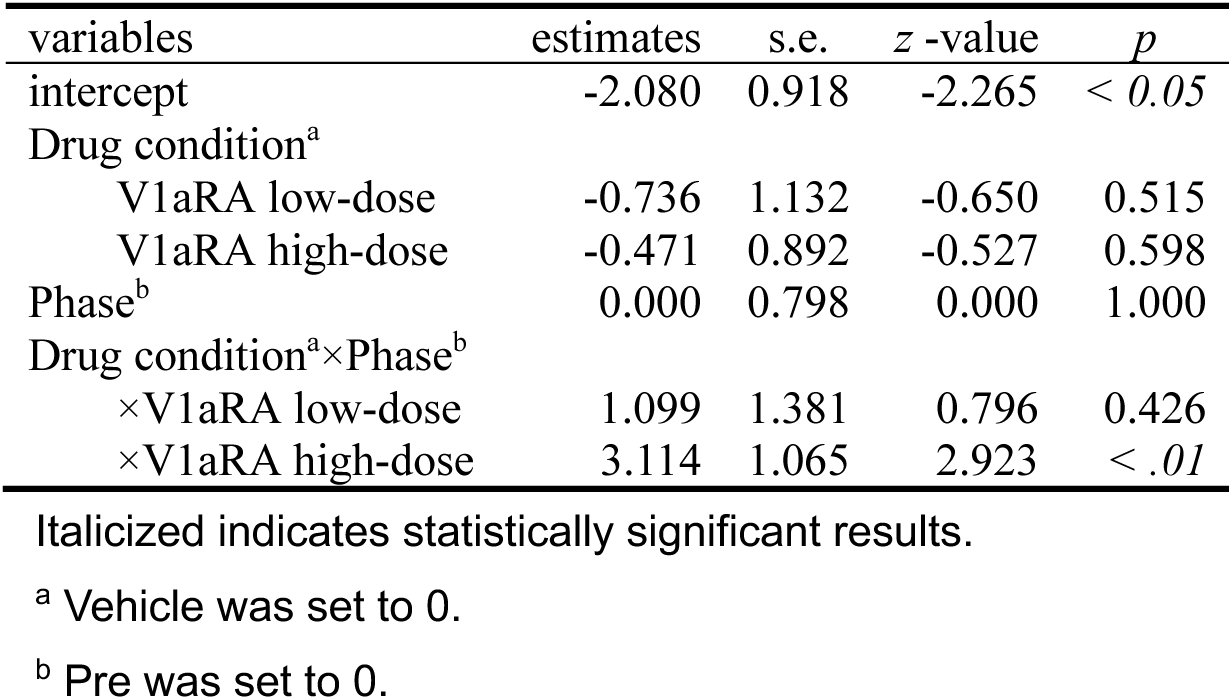
Outputs of GLMMs for aggressive behaviour in affiliated dyads.

| variables | estimates | s.e. | z -value | *p* |
| --- | --- | --- | --- | --- |
| intercept | -2.080 | 0.918 | -2.265 | < 0.05 |
| Drug condition<sup>a</sup> |  |  |  |  |
| V1aRA low-dose | -0.736 | 1.132 | -0.650 | 0.515 |
| V1aRA high-dose | -0.471 | 0.892 | -0.527 | 0.598 |
| Phase<sup>b</sup> | 0.000 | 0.798 | 0.000 | 1.000 |
| Drug condition<sup>a</sup>×Phase<sup>b</sup> |  |  |  |  |
| ×V1aRA low-dose | 1.099 | 1.381 | 0.796 | 0.426 |
| ×V1aRA high-dose | 3.114 | 1.065 | 2.923 | < .01 |

Italicized indicates statistically significant results.
<sup>a</sup> Vehicle was set to 0.
<sup>b</sup> Pre was set to 0.

**Table 7.** Post-hoc pairwise comparisons of pre- and post-administration on aggressive behaviour by drug conditions.

| Drug condition | Contrast | estimate | s.e. | z-ratio | *p* |
| --- | --- | --- | --- | --- | --- |
| Vehicle | Pre vs. Post | 0.000 | 0.798 | 0.000 | 1.000 |
| V1aRA low-dose | Pre vs. Post | -1.100 | 1.127 | -0.975 | 0.330 |
| V1aRA high-dose | Pre vs. Post | -3.110 | 0.705 | -4.416 | *< 0.001* |

*Italicized indicates statistically significant results.*

**Table 8.** Post-hoc pairwise comparisons of drug conditions on aggressive behaviour in the post-administration phase.

| Contrast | estimate | s.e. | z-ratio | *p* |
| --- | --- | --- | --- | --- |
| Vehicle vs. V1aRA low-dose | -0.362 | 0.805 | -0.450 | 0.894 |
| Vehicle vs. V1aRA high-dose | -2.643 | 0.584 | -4.523 | < 0.001 |
| V1aRA low-dose vs. V1aRA high-dose | -2.281 | 0.590 | -3.867 | < 0.001 |

*Italicized indicates statistically significant results.*

**Table 9.** Outputs of GLMMs for aggressive behaviour in unaffiliated dyads.

| variables | estimates | s.e. | z -value | *p* |
| --- | --- | --- | --- | --- |
| intercept | 0.182 | 0.289 | 0.632 | 0.528 |
| Drug condition<sup>a</sup> |  |  |  |  |
| V1aRA low-dose | 0.105 | 0.456 | 0.231 | 0.817 |
| V1aRA high-dose | -0.057 | 0.377 | -0.152 | 0.880 |
| Phase<sup>b</sup> | -0.087 | 0.417 | -0.208 | 0.835 |
| Drug condition<sup>a</sup>×Phase<sup>b</sup> |  |  |  |  |
| ×V1aRA low-dose | -0.047 | 0.665 | -0.070 | 0.944 |
| ×V1aRA high-dose | 0.198 | 0.535 | 0.371 | 0.711 |

*Italicized indicates statistically significant results.*
<sup>a</sup> Vehicle was set to 0.
<sup>b</sup> Pre was set to 0.

The administration of a high dose of the V1aR antagonist increased submissive vocalizations from subordinate individuals toward dominants but did not enhance them from dominants toward subordinates in affiliated dyads (Figure 6, Figure 4-6—source data 1, Video 2). GLMM analysis revealed a significant interaction between drug condition and phase under the high-dose V1aR antagonist condition (estimate ± s.e. = 2.036 ± 0.451, *p* < 0.001; Table 10). Post-hoc pairwise comparisons further clarified that submissive vocalizations in affiliated dyads significantly increased during the post-administration phase under the high-dose condition of the V1aR antagonist (*Pre vs. Post*, estimate ± s.e. = -2.110 ± 0.236, *p* < 0.001; Table 11). Moreover, post-hoc comparisons of drug conditions on submissive vocalizations in affiliated dyads during the post-administration phase revealed that the high-dose condition significantly increased submission compared to both the vehicle and low-dose conditions (*Vehicle vs. V1aRA high-dose*, estimate ± s.e. = -2.149 ± 0.279, *p* < 0.001; *V1aRA low-dose vs. V1aRA high-dose*, estimate ± s.e. = -2.538 ± 0.346, *p* < 0.001; Table 12). These findings indicate a substantial increase in submission from subordinate individuals following the administration of a high dose of the V1aR antagonist. No similar increase in submission was observed in dominant individuals (see Figure 6; GLMM analysis was not conducted due to the absence of submission under any condition or in any bird). Furthermore, the V1aR antagonist did not enhance submission in unaffiliated dyads (Figure 6 and Table 13).

**Figure 6.**
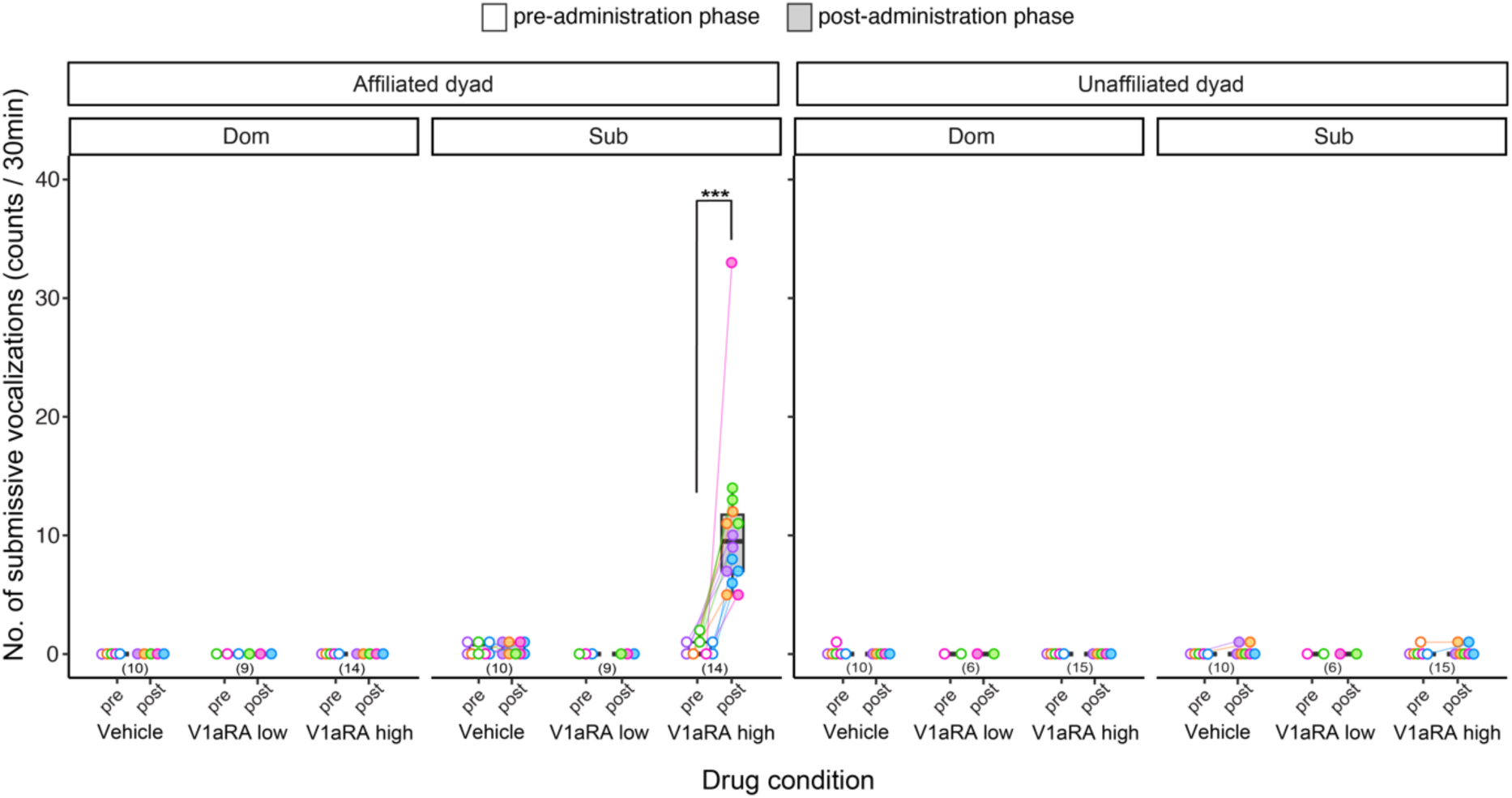
Effects of V1aRA on the number of submissive begging vocalizations in affiliated and unaffiliated dyads. In affiliated dyads, submissive vocalization initiated by the subordinate individual (SUB, right panel) exhibited a significant increase following administration of the *high-dose* V1aRA, as compared to pre-administration phase. In contrast, no such increase was observed in the low-dose or vehicle conditions. Additionally, this elevation in submissive vocalization was not observed in any condition for the dominant individual (DOM, left panel). Furthermore, in unaffiliated dyads, no significant effects were detected in either the dominant or subordinate individuals under any of the administered conditions. The plots illustrate individual data for each trial. The upper, middle, and lower bounds of the boxes represent the 75^th^, 50^th^, and 25^th^ percentiles, respectively, with whiskers extending to the maximum and minimum values. Numbers in the brackets below the boxes denote the number of samples (no. of birds × no. of trials). Asterisks denote statistically significant differences (Wald *χ^2^*statistics, *** *p* < 0.001). The colours correspond to the affiliated dyads shown in Figure 3.

**Table 10.**
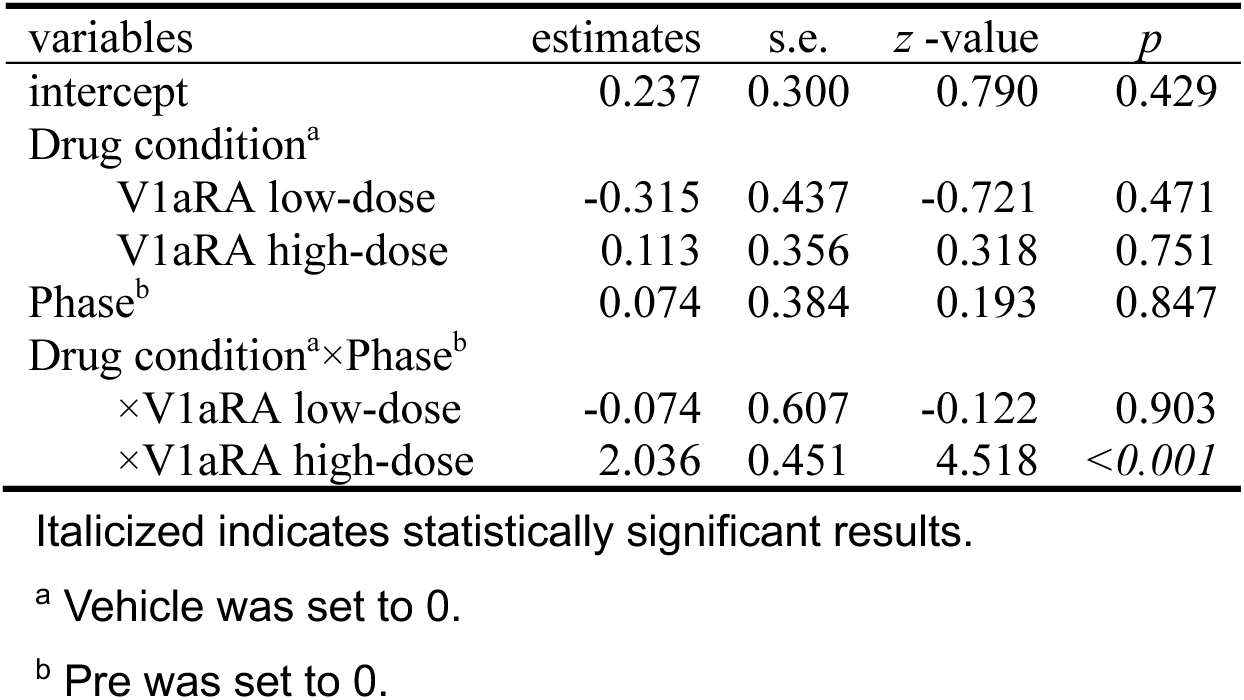
Outputs of GLMMs for submissive vocalization in affiliated dyads.

**Table 11.** Post-hoc pairwise comparisons of pre- and post-administration on submissive vocalization by drug conditions.

| Drug condition | Contrast | estimate | s.e. | z-ratio | *p* |
| --- | --- | --- | --- | --- | --- |
| Vehicle | Pre vs. Post | -0.074 | 0.384 | -0.193 | 0.847 |
| V1aRA low-dose | Pre vs. Post | 0.000 | 0.470 | 0.000 | 1.000 |
| V1aRA high-dose | Pre vs. Post | -2.110 | 0.236 | -8.942 | < 0.001 |

Italicized indicates statistically significant results.

**Table 12.** Post-hoc pairwise comparisons of drug conditions on submissive vocalization in the post-administration phase.

| Contrast | estimate | s.e. | z-ratio | *p* |
| --- | --- | --- | --- | --- |
| Vehicle vs. V1aRA low-dose | 0.389 | 0.431 | 0.903 | 0.638 |
| Vehicle vs. V1aRA high-dose | -2.149 | 0.279 | -7.699 | < 0.001 |
| V1aRA low-dose vs. V1aRA high-dose | -2.538 | 0.346 | -7.336 | < 0.001 |

Italicized indicates statistically significant results.

**Table 13.** Outputs of GLMMs for submissive vocalizations in unaffiliated dyads.

| variables | estimates | s.e. | z -value | *p* |
| --- | --- | --- | --- | --- |
| intercept | 0.000 | 0.316 | 0.000 | 1.000 |
| Drug condition<sup>a</sup> |  |  |  |  |
| V1aRA low-dose | 0.000 | 0.516 | 0.000 | 1.000 |
| V1aRA high-dose | 0.125 | 0.399 | 0.314 | 0.753 |
| Phase<sup>b</sup> | 0.182 | 0.428 | 0.426 | 0.670 |
| Drug condition<sup>a</sup>×Phase<sup>b</sup> |  |  |  |  |
| ×V1aRA low-dose | -0.182 | 0.719 | -0.254 | 0.800 |
| ×V1aRA high-dose | -0.125 | 0.546 | -0.229 | 0.819 |

Italicized indicates statistically significant results.
<sup>a</sup> Vehicle was set to 0.
<sup>b</sup> Pre was set to 0.

## Discussion

The present study suggested that in Experiment 1, triadic cohabitation facilitated the formation of an affiliative relationship exclusively between specific dominant and subordinate male crows within each triad. In Experiment 2, the peripheral administration of a V1aR antagonist disrupted these affiliative relationships, leading to a resurgence of aggressive and submissive/avoidant behaviours depending on the relative dominance but had no effect on social behaviours within unaffiliated dyads. Our findings suggest that the triadically interactive environment facilitative the formation of affiliative relationships between dominant and subordinate males. Moreover, the V1aR, previously implicated in same-sex affiliative relationships in studies on voles and finches, is essential in maintaining these types of male-male affiliative relationships.

### Experiment 1: The triadic cohabitation facilitated the formation of an affiliative relationship between two particular males

The results of Experiment 1 suggest that a triadically interactive situation plays a crucial role in facilitating the formation of affiliative relationships exclusively between two specific dominant and subordinate males of the three. Allopreening behaviour rarely occurred the dyadic cohabitation (i.e., pre-triadic cohabitation) but did frequently and exclusively between two specific males during triadic cohabitation. This finding suggests that triadic interactions are crucial for facilitating allopreening and the formation of affiliative relationships. Our finding aligns with previous theoretical studies (Caplow, 1956; Mesterton-Gibbons et al., 2011; Koykka & Wild, 2017), indicating that triad, including individuals with different power dynamics, are the minimal social groups that promote the formation of coalitions in the form of two against one.

This two-vs.-one form has been confirmed in previous behavioural observation studies on same-sex coalitions and alliances of primates, cetaceans, and corvids in the wild and in captivity (Macfarlan et al., 2014; von Rueden et al., 2008; von Rueden & Jaeggi, 2016; Patton, 2005; Watts, 2002; Muller & Mitani, 2005; Gilby et al., 2013; Bercovitch, 1988; Noë, 1994; Silk, 1999; Kutsukake & Hasegawa, 2005; Connor et al., 1992a; Connor et al., 1992b; Connor et al., 1999; Green et al., 2015; Kobayashi et al., 2020; Bygott et al., 1979; Caro, 1993; Caro, 1994; Fraser & Bugnyar, 2012). Our findings in Experiment 1 provide experimental evidence that a triadically interactive situation, such as 2-week cohabitation, facilitates the formation of affiliative relationships between dominant and subordinate males, bridging the gap between previous theoretical and behavioural observation studies.

The function of male-male affiliative relationships formed in Experiment 2 remains to be elucidated, but several possibilities warrant discussion. Generally, having affiliative partners may be beneficial in various ways. For large birds like crows, where predation pressure is relatively low (Braun et al., 2012), these relationships may primarily enhance foraging efficiency (Beauchamp, 2010; Dall & Wright, 2009). Experiments with captive *Corvus* species show that the presence of bonded individuals affects some behaviours, such as exploring new objects (Stöwe et al., 2006), paying attention to others (Scheid et al., 2007) and social learning (Schwab et al., 2008). Proximately, affiliation may serve as a social manoeuvre facilitating resource access and may assess the quality of a partner in these long-term monogamous birds (Braun & Bugnyar, 2012).

Another possibility is that crows use such affiliative interactions to maintain and intensify valuable social relationships, enabling coalition formation (de Waal & Luttrell, 1988). Studies in other *Corvus* species have shown a significant correlation between preening and agonistic support, suggesting that partners who engage in mutual preening are more likely to support each other in aggressive interactions (Scheid et al., 2007). Such behaviour can increase social rank for both individuals involved (Emery et al., 2007b). Common ravens focus on a few close partners at a time, aligning with the selective affiliative relationship formation observed in this study. Although agonistic support and subsequent rank reversals were not observed, this does not entirely rule out the possibility of coalition functions. Although agonistic support and subsequent rank reversals were not observed in our study, this does not entirely rule out the possibility of coalition functions. Aggressive support is much rarer than preening or grooming, making the opportunity to reciprocate support within a day very low. Ravens have been shown to reciprocate support over two years rather than within a week, reflecting the likelihood of long-term service exchange (Fraser & Bugnyar, 2012). Thus, coalition-like behaviours might not have been observed due to our study’s specific context and observation period but might be evident over longer time spans or different contexts. Future experiments could test the coalition function in male-male affiliative relationships by confronting affiliated dyads formed in triadic cohabitation with a male-female pair or by introducing them into a captive group to observe rank changes. The high levels of affiliative interactions maintained during post-dyadic cohabitation suggest potential for long-term alliances. Long-term collaborations in conflicts, referred to as alliances, contrast with facultative short-term coalitions (Harcourt & de Waal, 1992). Our study implies that the affiliative relationships formed in triadic cohabitation may constitute alliances, warranting further investigation into their stability when a third-party individual is introduced or when the dyad is integrated into larger groups.

Male-male affiliative relationships in our crows may also function as social bonds similar to those seen in mother-offspring or pair bonds, potentially reducing social stress. The presence of affiliative individuals mitigates stress responses, a phenomenon known as social buffering (Kikusui et al., 2006). Many animals, including humans, exhibit smaller increases in stress hormone (cortisol) levels when exposed to aversive stimuli in the presence of an affiliative partner compared to when alone (de Vries et al., 2003). In primates, social grooming reduces heart rate (Aureli et al., 1999), and individuals with tightly-knit social networks have lower baseline levels of cortisol metabolites in their faeces (Brent et al., 2011; Crockford et al., 2008). Following concepts developed in primatology (Hinde, 1976), social bonds in Corvids are defined by the reciprocal exchange of affiliative behaviours over time, observed across all age classes and between both male-female and male-male dyads (Braun et al., 2012). In ravens, individuals with high levels of positive social connections within their group show lower corticosterone levels when within the group and higher levels when separated (Stocker et al., 2016). Affiliated dyads in our study might have reduced stress from attacks by the dominant and the stress of living in confined cages through mutual preening. In large-billed crows, methods for measuring corticosterone in both faeces and saliva using enzyme-linked immunosorbent assay have been established (Ode et al., 2015; Aota et al., 2023). Future studies should compare corticosterone levels in saliva or faeces during both the early and last periods of triadic cohabitation to determine whether a decrease in corticosterone accompanies the formation of affiliative relationships. Additionally, examining changes in corticosterone levels during separation from and reunion with affiliative partners would provide further insights into the stress-buffering role of these relationships.

While numerous challenges remain in understanding the functions of male-male affiliative relationships in large-billed crows, the hypotheses discussed above are not mutually exclusive. Applying triadic cohabitation experiments as described can provide valuable insights through careful and systematic testing. Moreover, these experiments can help to identify predictors of affiliative relationship formation among males. Several studies have concluded that personality similarity or homophily is a strong predictor of affiliative relationships, potentially facilitating cooperation (Verspeek et al., 2019; Laakasuo et al., 2020; Noë et al., 2016; Massen & Koski, 2014). By collecting data on inter-individual behavioural differences or personalities, such as exploration tendencies and stress vulnerability before triadic cohabitation, we can examine which characteristics, beyond social rank, kinship or age, predict affiliative relationships. Triadic cohabitation experiments enable physiological exploration and rigorous testing of predictors, enhancing our understanding of the mechanisms underlying affiliative relationship formation.

Two of the eight triads tested in the present study did not form affiliative relationships even after 2-week cohabitation. As illustrated in Figure 3, allopreening occurred slightly in two males of the two triads during the cohabitation period but eventually disappeared. According to the video observation of behaviours during the cohabitation period, one possible cause for the failure to form affiliative relationships in these triads is that one male avoided the initiation of allopreening by the other male. Another possible cause is the intervention by the third-party male when the two males attempted to engage in socially affiliative interactions. The intervention by third-party individual in affiliative interactions between other individuals has been found in a captive group of ravens (Massen et al., 2014b). The questions of what social characteristics of individuals in dyads could facilitate or inhibit allopreening and what individual and/or social factors could drive the intervention in others’ affiliative behaviour in birds and mammals remain to be addressed. An in-depth analysis of social behaviour using the triadic cohabitation paradigm used in this study may elucidate the behavioural and psychological mechanisms for forming male-male affiliative relationships.

### V1a receptor antagonism disrupted the dominant-subordinate affiliative relationship

Systemic injection of a V1aR antagonist to males with affiliative relationships caused a decrease in allopreening and the resurgence of agonistic behaviours associated with relative dominance, such as aggressive behaviour in the dominant and submissive behaviour (i.e., submissive begging calls) in the subordinate. In contrast, dominant and subordinate males without an affiliated relationship showed no behavioural effects from V1aR blocking. These results suggest that V1aR is involved in the suppression of dominance-dependent aggressive/submissive behaviour and facilitating affiliative behaviour, thus playing a crucial role in the formation and maintenance of affiliated relationships between dominant and subordinate males.

The dose-dependent effects of the V1aR antagonist on allopreening, aggression, and submission raise important points for discussion. The finding that allopreening significantly decreased even at low doses, while aggression and submission showed significant changes only at high doses, suggests that suppressing antagonistic behaviours may be a prerequisite for the occurrence of allopreening, with V1a receptors playing a key role. Indeed, as observed in Experiment 1 of this study, aggression and submission between two individuals first decreased (Figure 2), followed by an increase in preening during the triadic cohabitation period (Figure 3A). It has also been proposed that affiliative behaviours progress hierarchically, from tolerance of proximity to more intimate interactions, such as grooming and food sharing (Silk et al., 2013). Further research is required to investigate the relationship between the depth of social relationships and dose-dependent effects across species.

Given the peripheral administration of the drug in this study, the behavioural effects found in affiliated dyads from V1aR antagonists do not necessarily reflect the role of V1aR in the brain. Peripheral nonapeptide receptors have been suggested to influence a variety of homeostatic functions, such as hydromineral balance (Goldstein, 2006) and thermoregulation (Hassinen et al., 1999). For example, maternal crouching in rodents serves a thermoregulatory function (Jans & Woodside, 1990) and, is regulated by V1aR (Bosch & Neumann, 2008).

V1aR has also been suggested to be involved in peripheral thermoregulation (Pittman et al., 1998; Shido et al., 1984; Yang et al., 2006). Thus, the effects of maternal crouching by V1aR blocking in rodents may be indirectly due to thermoregulation via V1aR antagonism. Similarly, maternal licking and grooming in rodents may indirectly involve peripheral V1aR (Bosch & Neumann, 2008). Given that during maternal licking and grooming, mothers often ingest water to maintain the hydromineral balance (Gubernick & Alberts, 1983), based on peripheral V1aR (Koshimizu et al., 2012), maternal licking and grooming may be regulated indirectly by peripheral V1aR. Although the distribution of V1aR (i.e., VT4R) has been examined in the brain and pituitary gland (Leung et al., 2009; Selvam et al., 2013) but not in peripheral organs in birds, previous studies reporting the involvement of VT in renal osmoregulation (Goldstein, 2006; Braun & Dantzler, 1984) suggest the presence of V1aR in the periphery of birds. Thus, we cannot exclude the possibility that peripheral injection of a V1aR antagonist affected osmoregulation and secondary social behaviour effects. Although we did not measure the effects of body temperature and the amount of drinking/evacuation following V1aR antagonist injection, such homeostatic effects on the periphery from V1aR blocking are unlikely to occur selectively in affiliated dyads. To determine the involvement of the central V1aR in the social behaviour of affiliated males, it is necessary to examine the effect of central infusion of V1aR antagonists in future studies.

### Differences in functions and inherent aggression in same-sex affiliative relationships: a comparison of crows and other species

There are various types of same-sex affiliative relationships that differ in sex, kinship and function. The underlying physiological mechanisms might also be diverse. The extensive body of research on meadow voles (Anacker et al., 2016) and zebra finches (Goodson & Wang, 2006; Kelly & Goodson, 2013; Kelly et al., 2011) has established that V1aR plays a crucial role in promoting same-sex affiliative relationships or behaviours. In this study on large-billed crows, it was revealed that V1aR is involved, although the function of same-sex affiliation and the inherent agonistic tendencies between individuals differ from those in voles and finches.

Meadow voles and prairie voles exhibit a behavioural preference for familiar individuals over unfamiliar ones, engaging in affiliative interactions and forming communal groups with minimal aggression (Madison & McShea, 1987; Getz et al., 1993). Both voles share common functions in maintaining body temperature and predator defence through group living, which supports safe breeding (Madison & McShea, 1987; Getz et al., 1993). Meadow voles prefer familiar individuals during winter in the wild and under short day winter-like conditions in the lab, huddling together. Winter social groups, seeded by a female and her undispersed offspring (mother-daughter or sister-sister relationships), often continue into spring with communal nesting (McShea & Madison, 1984).

Prairie voles, closely related to meadow voles, have been primarily studied for male-female interactions, but same-sex interactions have also been investigated. Regardless of day length, both males and females prefer familiar mates and same-sex peers (DeVries et al., 1997; Beery et al., 2018; Lee et al., 2019). This preference, without seasonal variation and not biased towards females, differs from that of meadow voles. However, prairie vole units are fundamentally composed of bonded breeding pairs and undispersed offspring (Getz et al., 1993; Carter & Getz, 1993). These findings suggest that same-sex affiliative relationships in both species of voles provide not only thermoregulatory benefits, as commonly demonstrated in rodents (Andrews et al., 1987; Andrews & Belknap, 1993; Gilbert et al., 2010; Kauffman et al., 2003), but also ensure the safe rearing of offspring.

Gregarious zebra finches are highly social and tolerant, roosting and breeding together in colonies for anti-predator defense (Zann, 1996; Schuett & Dall, 2010; Evans et al., 2018), with low-intensity aggression such as threats and chases (Ruploh et al., 2013). Gregariousness is a term used to describe species that affiliate with or live in groups (Miller, 1922; Treisman, 1975; Goodson & Kingsbury, 2011; Peterson & Weckerly, 2018). Recent studies of neuropeptide-mediated behaviours have also focused on spiny mice (*Acomys cahirinus*), group-living communal breeders, which also exhibit a gregarious phenotype (Kelly & Seifert, 2021; Powell et al., 2023; Fricker & Kelly., 2024; Fricker et al., 2024). In spiny mice, both males and females show little aggression in various social contexts and prefer affiliating with larger groups (Fricker et al., 2023; Gonzalez Abreu et al., 2022; Fricker et al., 2024).

Research on voles, finches, and spiny mice has shed light on the physiological basis of same-sex affiliative relationships, including the role of V1aR. However, since these species exhibit low levels of inter-individual aggression, the exploration of the physiological basis of affiliative relationships formed between dominant and subordinate individuals has not been investigated. As previously mentioned, the exact function of the affiliative relationships formed between dominant and subordinate males in large-billed crows has not yet been clarified. However, in other *Corvus* species, these relationships are known to function as coalitions and provide social buffering effects (Fraser & Bugnyar, 2012; Braun & Bugnyar, 2012; Fraser & Bugnyar, 2010b; Fraser & Bygnyar, 2011; Seed et al., 2007). Therefore, it is highly likely that similar functions exist in our study subjects as well. The involvement of V1aR demonstrated in this study of male-male affiliative relationships suggests a universal role for V1aR in same-sex affiliative relationships despite the likely fundamental differences in function and the levels of inherent aggression between individuals compared to voles and finches.

One of our most intriguing findings was that the V1aR antagonist disrupted the affiliative relationship, causing the resurgence of aggressive and submissive behaviours depending on the relative dominance only within affiliated dyads but had no effect within unaffiliated dyads. Given this result, the role of V1aR in male-male affiliation among crows might be to inhibit the inherent agonistic tendencies and enable affiliative interactions, such as allopreening and close proximity. This function of V1aR might have been evolutionarily acquired in crows and, possibly in primates and dolphins as well. Similar context-specific involvement of V1aR in the control of aggressive behaviour in birds has been suggested in previous studies. For instance, a previous study on violet-eared waxbills (*Uraeginthus granatina*) found that peripheral injection of a V1aR antagonist disinhibited and increased the aggression of subordinate males in a territorial challenge context, whereas V1aR reduced the aggression of dominant males towards rival males in a mate competition context (Goodson et al., 2009). However, in their study, male-male interactions were highly antagonistic and did not form affiliative relationships, making it impossible to verify the connection between affiliation and dominance relationships. To clarify the role of V1aR and the relevant VP/VT signals in facilitating affiliative behaviour and/or inhibiting agonistic behaviour in affiliated males, future studies should investigate the central infusion of a V1aR antagonist and assess V1aR-/VT-neuron activities using crows. Additionally, recent studies on localisation of V1aR and OTR in chimpanzees and other primates (Rogers Flattery et al., 2022) suggest promising developments in this field.

In conclusion, we demonstrated that a triadically interactive environment plays a crucial role in forming male-male affiliative relationships in non-human animals. Our study suggests that dominant-subordinate affiliative relationships in males share common neural underpinnings with other same-sex bonds, pair bonds and parent-infant bonds involving the VP/VT systems. This finding provides insights into social affiliation, indicating that VP/VT may facilitate these relationships by promoting affiliative behaviours and suppressing aggression. Our results underscore the significance of triadic interactions in social behaviour, providing a new perspective on how complex social relationships are formed and sustained. Future research should explore the roles of VP/VT as well as OT/MT and their receptor subtypes in social affiliation among same-sex individuals to enhance our understanding of the evolution of inter-individual affiliation in the social lives of various species, including humans.

## Acknowledgements

We thank the members of the Animal Psychology Lab, Keio University, for their inspiring scientific discussions and their support in the daily care of the animals. We also thank Dr Shigeru Watanabe (Keio University, Japan) for his helpful advice and discussions. This work was financially supported by JSPS KAKENHI (#23H01058, #23H00074, #23K17645, #20H01787 to E.-I., and #19J22654 to A.S.) and a Keio University Grant-in-Aid for ICRP (#MKJ1905) to E.-I.

## Author contributions

A.S. designed research, conducted the experiments, analysed data, wrote the manuscript, and acquired funding. E.-I. reviewed and edited the manuscript, supervised the work, and acquired funding.

## Declaration of interests

The authors declare no competing interests.

## Source data and videos

Source data and videos related to Experiments 1 and 2 can be found online at https://figshare.com/projects/Seguchi_Izawa_Role_of_Vasopressin1a_on_male-male_affiliations_in_large-billed_crows/213883

Figure 2—source data 1

: Source data of social behaviours in dyadic encounters before Experiment 1 (Figure 2). Figure 3—source data 1

: Source data of allopreening behaviours in Experiment 1 (Figure 3A). Figure 3—source data 2

: Source data of allopreening behaviours used in discriminant analysis (Figure 3B). Figure 4-6—source data 1

: Source data of social behaviours in Experiment 2 (Figure 4-6). Video 1

: Triadic cohabitation. Video 2

: Effects of V1aR antagonism.

## Materials and Methods

### Key resources table

| REAGENT or<br>RESOURCE | SOURCE | IDENTIFIER |
| --- | --- | --- |
| Chemicals |  |  |
| SR49059 | Sigma-Aldrich | CAS# 150375-75-0 |
| Dimethyl sulfoxide | Fujifilm | CAS# 67-68-5 |
| Deposited data |  |  |
| Data for figures and<br>tables | This paper | https://figshare.com/projects/Seguchi_Izawa_Role_of_Vasopressin1a_on_male-male_affiliations_in_large-billed_crows/213883 |
| Software |  |  |
| SPSS | IBM Corp. (2021). IBM SPSS<br>Statistics for Windows, Version<br>28.0. IBM Corp, Armonk, NY,<br>USA. URL<br>https://www.ibm.com/products/spss-statistics. | NA |
| R, RStudio | RStudio Team (2020). RStudio:<br>Integrated Development for R.<br>RStudio, PBC, Boston, MA,<br>USA. URL http://www.rstudio.com/. | NA |

### Experimental Model and Subject Details

We used 18 adult male crows aged between 4 and 7 years, with body weights ranging from 640 to 790 g. Sex was determined using DNA from blood samples (Fridolfsson & Ellegren, 1999). The birds were caught as yearlings in Tokyo and neighbouring areas during 2017–2019, with permission from the Ministry of the Environment Government of Japan (authorisation nos. 29030001, 30081482, 30030019, and 30224001). After capture, the crows were housed in one of eight different groups with 5–10 other crows (Table 14) in outdoor aviaries (W 5 m × D 10 m × H 3 m). Three to six months before the start of this study, the crows were transported to individual steel-mesh cages (W 57 cm × D 93 cm × H 63 cm) in an animal housing room to prevent the formation of dominance or affiliative relationships through physical interactions. Individual cages were placed side by side on shelves in the room, allowing the birds to see and hear each other but preventing physical contact to reduce physical and social stress due to individual housing. Room temperature was maintained at 21 ± 2 ℃, with a light/dark cycle of 13 hours light and 11 hours dark, starting at 0800. Food (dried dog food and other supplements) and water were freely available in the cages.

**Table 14.** Composition of the subjects in the group housing prior to this study and their ages.

| *ID* | *Group* | *Age* |
| --- | --- | --- |
| VV | Group A | 6 |
| RR |  | 6 |
| OW |  | 6 |
| PY |  | 6 |
| BB |  | 6 |
| #28 | Group B | 6 |
| #27 |  | 6 |
| KR | Group C | 6 |
| KY |  | 6 |
| KV |  | 6 |
| KG |  | 6 |
| LK | Group D | 4 |
| GK |  | 4 |
| YA | Group E | 7 |
| OBO | Group F | 4 |
| RM | Group G | 7 |
| VA | Group H | 6 |

Letters indicate individual birds.

The experiments were conducted from October 2019 to September 2022. Due to the accidental death of some birds and the restricted use of the experimental facility during the COVID-19 pandemic, some of the planned experiments were not performed, resulting in unbalanced data. The experimental and housing protocols adhered to the Japanese National Regulations for Animal Welfare and were approved by the Animal Care and Use Committee of Keio University (#20036, #A2022-055).

## Method Details

### Determination of relative dominance in dyads

We assigned 18 birds to 8 triads for the cohabitation experiment (Table 1). Before assigning the triad groups, we determined the relative dominance between each pair of birds through three trials of dyadic social encounters in an indoor aviary (W 2 m × D 2 m × H 2 m), using a procedure similar to that reported in previous studies (Nishizawa et al., 2011; Takeda et al., 2022). In each trial, two birds were given a 5-min opportunity to interact freely. The winner and loser were determined according to the behavioural criteria used in previous studies (Nishizawa et al., 2011; Takeda et al., 2022).

Specifically, the loser was identified as the bird exhibiting submissive behaviour (i.e., submissive begging vocalisation and/or avoidance) in response to aggressive behaviour (i.e., jab, peck, aggressive vocalisation and approach). Within the dyad, the bird who won all three trials was defined as dominant, while the other as subordinate. All 26 dyads showed an apparent asymmetry in win/loss numbers across the three trials, indicating the formation of dominance relationships (Figure 2A, B, Figure 2—source data 1). Thus, 26 dyads with clear dominance relationships were used in Experiment 1. The social encounter trials were conducted between 10:00 and 15:00, with a 2-day interval between consecutive trials involving the same dyads. No subjects showed allopreening in any trial, indicating that no affiliative relationship was formed in any dyad at this stage (Figure 2C, Figure 2—source data 1).

### Experiment 1: The triadic cohabitation facilitated the formation of an affiliative relationship between two particular males

After determining the relative dominance positions of the individuals in the dyads of each triad, we tested whether a 2-week cohabitation in triads facilitated the formation of affiliative relationships between two specific birds in the triad (Figure 1A). The formation of an affiliative relationship was evaluated based on the emergence and increase in reciprocal allopreening exclusively between two specific birds in each triad. To confirm that affiliative relationships were not formed through dyadic interactions, all possible dyads of each triad (i.e., three dyads) were housed together in the experimental aviary on separate occasions for three days before triadic cohabitation (hereafter referred to as pre-test). This pre-test phase was followed by the 2-week triadic cohabitation to examine whether triadic interactions could facilitate the formation of affiliative relationships. To ensure that the relationships observed during the triadic cohabitation were not simply temporary or artefacts of spatial arrangements, a second dyadic cohabitation period was conducted following the triadic cohabitation (hereafter referred to as post-test). In the pre- and post-tests, each of the two birds was transferred from its individual home cage into the aviary at 8:00 am, and after 72 h, it was returned to the home cage at 8:00 am. On the day following the end of the pre-test period, the three birds were introduced into the experimental aviary and housed together for two weeks. At the end of the triadic cohabitation, the birds were transferred back to their individual home cages. At 8:00 am the following day, each pair of the triad was again transferred into the aviary for 3 days as a post-test on separate occasions.

The behaviour of the birds during the pre-/post-tests and triadic cohabitation was video recorded using a ceiling-mounted camera (EX-ZR4100, CASIO, Japan) in the aviary. Recordings were conducted for 30 min randomly between 9:00 and 17:00 every day in the pre- and post-tests and for the early (1−3 days), middle (7−9 days), and last (12−14 days) three days of triadic cohabitation. Therefore, 90-min video data (30 min × 3 days) were used to analyse social interactions in each test, as well as in the 3 × 3-day blocks during the triadic cohabitation period. In the aviary, three wooden perches were placed in parallel with 30 cm intervals at a height of 80 cm above the floor. Food, water, and a bathing tub were consistently provided, except during the 30-min video recording sessions.

### Experiment 2: V1a receptor antagonism disrupted the dominant-subordinate affiliative relationship

To investigate the involvement of V1a-like receptors in the maintenance of male-male affiliative relationships, birds forming affiliated relationships in Experiment 1 were systematically administered a V1aR antagonist (SR 49059, Sigma-Aldrich, MO, USA) to examine the effects on social behaviours towards the affiliated counterpart and the unaffiliated male (Figure 1B and Table 1). This V1aR is an effective antagonist of the VT4 receptor in birds (Kuenzel et al., 2016). The effect of the V1aR antagonist was tested with high dose (0.3 mL × 1 mg/mL vehicle solution) and low dose (0.3 mL × 0.1 mg/mL vehicle solution), compared with a 0.3 mL vehicle solution as a control, administered via intramuscular injection into the chest of subject crows. The vehicle solution contained 10% dimethyl sulfoxide (Fujifilm, Tokyo, Japan) in a saline solution.

In each trial, the V1aR antagonist (high or low dose) or vehicle was administered to either the dominant or subordinate bird. Each bird was tested in eight trials: three with the high-dose condition, three with the low-dose condition, and two with the vehicle control condition. The trial order of the dominance position of the focal individual (dominant or subordinate) and the drug conditions were counterbalanced within and between dyads. After testing the affiliated dyads, the birds underwent three additional trials of dyadic interactions without drug injection to ensure the maintenance or recovery of the relationship. At least one week after this recovery trial, both birds of affiliated dyads received another set of eight trials with the unaffiliated individual (the third bird in the triadic cohabitation in Experiment 1) as an unaffiliated dyad test with the same procedure as that used in the affiliated dyad test.

For each trial, two crows were observed interacting in the aviary for 30 minutes in the morning (pre-administration phase). Following this interaction, either the antagonist or vehicle was administered to the dominant or subordinate bird (the focal bird) one hour prior to the next interaction. During this 1-hour isolation period, the focal bird was housed separately in an individual cage, isolated from the other bird. The trial began when the focal bird was introduced into the indoor aviary (W 2 m × D 2 m × H 2 m), with the other bird being introduced 1 minute earlier. Social behaviours, including allopreening, aggressive behaviours (such as peck, jab, and kick), and submissive begging vocalizations, were recorded from video data for offline analysis. The inter-trial interval ranged from three to five days, with each bird participating in one trial per day. All trials were conducted between 8:30 AM and 12:00 PM.

## Statistical Analysis

### Experiment 1: The triadic cohabitation facilitated the formation of an affiliative relationship between two particular males

We defined an affiliative relationship as a dyad where reciprocal, but not unidirectional, allopreening occurred between two birds, independent of their dominance relationship. This definition aligns the behavioural characteristics of affiliative relationships in primates and other social animals, such as maintaining close proximity and exchanging affiliative behaviours (Fedurek & Dunbar, 2009). An allopreening bout was counted when a bird passed its bill through the feathers of another bird for longer than 2 s (Miyazawa et al., 2020). Given the absence of allopreening in any dyad before triadic cohabitation, we defined a dyad as having an affiliative relationship if both individuals initiated allopreening in 10 or more bouts per 30 min in any of the 3-day blocks during the triadic cohabitation (Figure 3A, Figure 3—source data 1). Dyads not meeting this behavioural criterion were considered as not having formed an affiliative relationship. To confirm the exclusive formation of affiliative relationships through triadic cohabitation, we performed a discriminant analysis to determine whether the numbers of dominant-initiated and subordinate-initiated allopreening in the pre- and post-triadic cohabitation tests could distinguish affiliated dyads from their pre-relationship state and unaffiliated dyads (Figure 3B, Figure 3—source data 2). Leave-one-out cross-validation was used to calculate the correct discrimination rate. The discriminant analysis was performed using SPSS Statistics version 29. Allopreening behaviours were coded using BORIS v.7 (Frinard & Gamba, 2016).

### Experiment 2: V1a receptor antagonism disrupted the dominant-subordinate affiliative relationship

To evaluate the effects of subject dominance status, drug condition, and phase on social behaviour, we employed Generalized Linear Mixed Models (GLMMs) with a Poisson error distribution and a log link function. We analysed allopreening behaviours (Figure 4, Figure 4-6—source data 1 and Table 3-5), aggressive behaviours (Figure 5, Figure 4-6—source data 1Tables 6-9), and submissive vocalisations (Figure 6, Figure 4-6—source data 1Tables 10-13) separately. The model incorporated the number of behaviours per 30 min as the response variable, and three fixed factors: *subject dominance status* (dominant vs. subordinate), *drug condition* (vehicle, V1aRA low dose, V1aRA high dose), and *phase* (pre-administration vs. post-administration). Additionally, we examined the three-way interaction among these factors to determine whether the drug effects varied depending on dominance status and phase. *Trial number* and *identity of focal dyad* were included as random factors.

The significance of the fixed effects was assessed using Wald χ² statistics. If the interaction was significant, post-hoc pairwise comparisons were performed to further investigate significant interactions. Specifically, comparisons were made between the pre- and post-administration phases within each drug condition and dominance status, with results presented on the original scale.

Statistical significance was determined at the 5% level. All statistical analyses were performed using R version 4.4.1. The GLMMs were fitted using the glmer function from the package ‘lme4’ (Bates et al., 2015), and post-hoc pairwise comparisons were performed using the package ‘emmeans’(Lenth, 2023).

## Notes

https://figshare.com/projects/Seguchi_Izawa_Role_of_Vasopressin1a_on_male-male_affiliations_in_large-billed_crows/213883

### Competing Interest Statement

The authors have declared no competing interest.

### Summary of Updates

In the previous manuscript, the comparison between the vehicle data and the V1a receptor antagonist condition was simplistic, and we did not perform a comparison between pre- and post-administration for each drug condition. In the revised manuscript, we have included graphs and detailed descriptions comparing affiliative, aggressive, and submissive behaviors in male subjects before and after treatment under the high-dose V1aR antagonist, low-dose V1aR antagonist, and vehicle conditions. The conclusions that V1aR antagonists reduce allopreening between affiliated males and restore aggressive behaviour in dominant individuals and submissive vocalizations in subordinate individuals remain unchanged.

